# Integrating gene expression and imaging data across Visium capture areas with visiumStitched

**DOI:** 10.1101/2024.08.08.607222

**Authors:** Nicholas J. Eagles, Svitlana V. Bach, Madhavi Tippani, Prashanthi Ravichandran, Yufeng Du, Ryan A. Miller, Thomas M. Hyde, Stephanie C. Page, Keri Martinowich, Leonardo Collado-Torres

**Affiliations:** Lieber Institute for Brain Development, Johns Hopkins Medical Campus, 21205 Baltimore, USA; Department of Biomedical Engineering, Johns Hopkins School of Medicine, 21218, Baltimore, USA; Department of Psychiatry and Behavioral Sciences, Johns Hopkins School of Medicine, 21205, Baltimore, USA; Department of Neurology, Johns Hopkins School of Medicine, 21205, Baltimore, USA; The Solomon H. Snyder Department of Neuroscience, Johns Hopkins School of Medicine, 21205, Baltimore, USA; Johns Hopkins Kavli Neuroscience Discovery Institute, Johns Hopkins University, 21218, Baltimore, USA; Department of Biostatistics, Johns Hopkins Bloomberg School of Public Health, 21205, Baltimore, USA; Center for Computational Biology, Johns Hopkins University, 21205, Baltimore, USA

**Author notes:** Full list of author information is available at the end of the article.

**Keywords:** spatial transcriptomics, Visium, multi-capture area study design, image stitching

## Abstract

**Background:** Visium is a widely-used spatially-resolved transcriptomics assay available from 10x Genomics. Standard Visium capture areas (6.5mm by 6.5mm) limit the survey of larger tissue structures, but combining overlapping images and associated gene expression data allow for more complex study designs. Current software can handle nested or partial image overlaps, but is designed for merging up to two capture areas, and cannot account for some technical scenarios related to capture area alignment.

**Results:** We generated Visium data from a postmortem human tissue sample such that two capture areas were partially overlapping and a third one was adjacent. We developed the R/Bioconductor package *visiumStitched*, which facilitates stitching the images together with *Fiji* (*ImageJ*), and constructing *SpatialExperiment* R objects with the stitched images and gene expression data. *visiumStitched* constructs an artificial hexagonal array grid which allows seamless downstream analyses such as spatially-aware clustering without discarding data from overlapping spots. Data stitched with *visiumStitched* can then be interactively visualized with *spatialLIBD*.

**Conclusions:** *visiumStitched* provides a simple, but flexible framework to handle various multi-capture area study design scenarios. Specifically, it resolves a data processing step without disrupting analysis workflows and without discarding data from overlapping spots. *visiumStiched* relies on affine transformations by *Fiji*, which have limitations and are less accurate when aligning against an atlas or other situations. *visiumStiched* provides an easy-to-use solution which expands possibilities for designing multi-capture area study designs.

## Background

Among sequence-based spatially-resolved transcriptomics assays, the commercially-available Visium assay developed by 10x Genomics is one of the most commonly used assays [1, 2]. Visium v1 gene expression slides contain four capture areas, each of which can accommodate a tissue sample up to 6.5mm by 6.5mm in size. While other versions of Visium accommodate larger capture areas, they all ultimately have a fixed size and thus, there will be situations in which more than one capture area is required for specific research questions.

While Visium is a powerful tool for profiling spatially-resolved gene expression, capture areas for this platform are limited in size, and hence many tissue regions of interest can’t fit within a single capture area. One approach is to split the tissue of interest across multiple capture areas, and then treat data from each capture area as a separate sample for analysis; however, this approach discards spatial information about how the samples relate to one another, which is important for biological context. To address this problem, several approaches have been suggested [3, 4, 5, 6, 7]. However, existing solutions either do not address both imaging and genomics data simultaneously, cannot handle partial overlap of capture areas, cannot directly be applied to samples consisting of three or more capture areas, or do not facilitate using the stitched data in downstream analyses via a new artificial Visium array (**Table 1**).

**Table 1.**
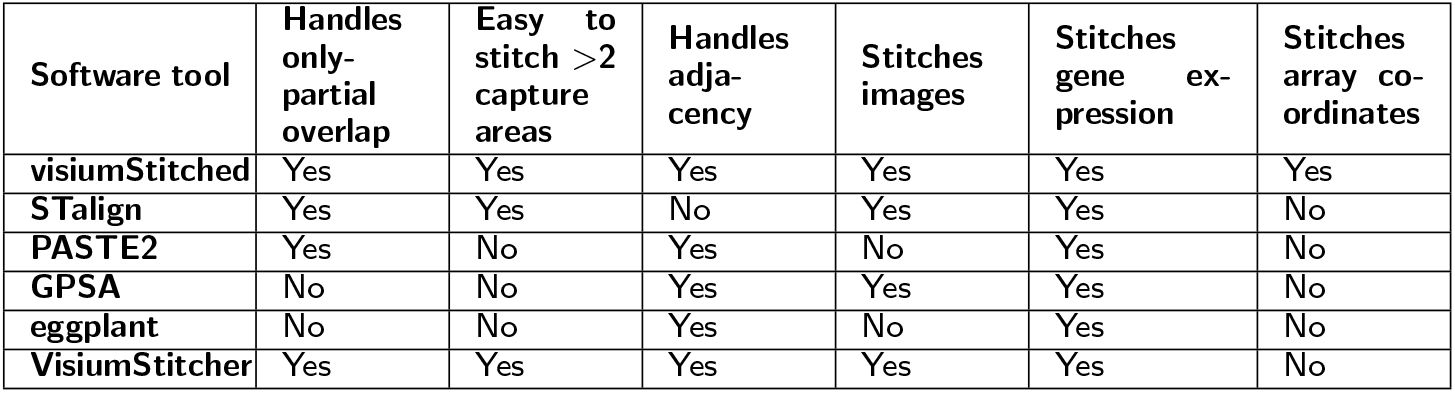
Comparison of software tools for stitching spatial transcriptomics data. Many existing tools are unable to handle adjacent or partially overlapping capture areas, do not stitch both image and gene-expression data, or do not directly support the stitching of more than two capture areas at once.

We introduce *visiumStitched*, an R/Bioconductor package building upon the functionality of *Fiji* [8], to stitch imaging and gene-expression data from separate capture areas into coherent, analysis-ready R/Bioconductor *SpatialExperiment* objects [9]. *visiumStitched* is designed to allow researchers to treat the entire tissue region of interest as a single Visium sample, rather than performing analyses separately on individual Visium capture areas. *SpatialExperiment* objects created with *visiumStitched* are compatible with downstream spatial clustering methods such as *BayesSpace* and *PRECAST* [10, 11]. To demonstrate the utility of *visiumStitched*, we generated sample Visium data across three capture areas from a postmortem human brain donor (**Figure 1**).

**Figure 1.**
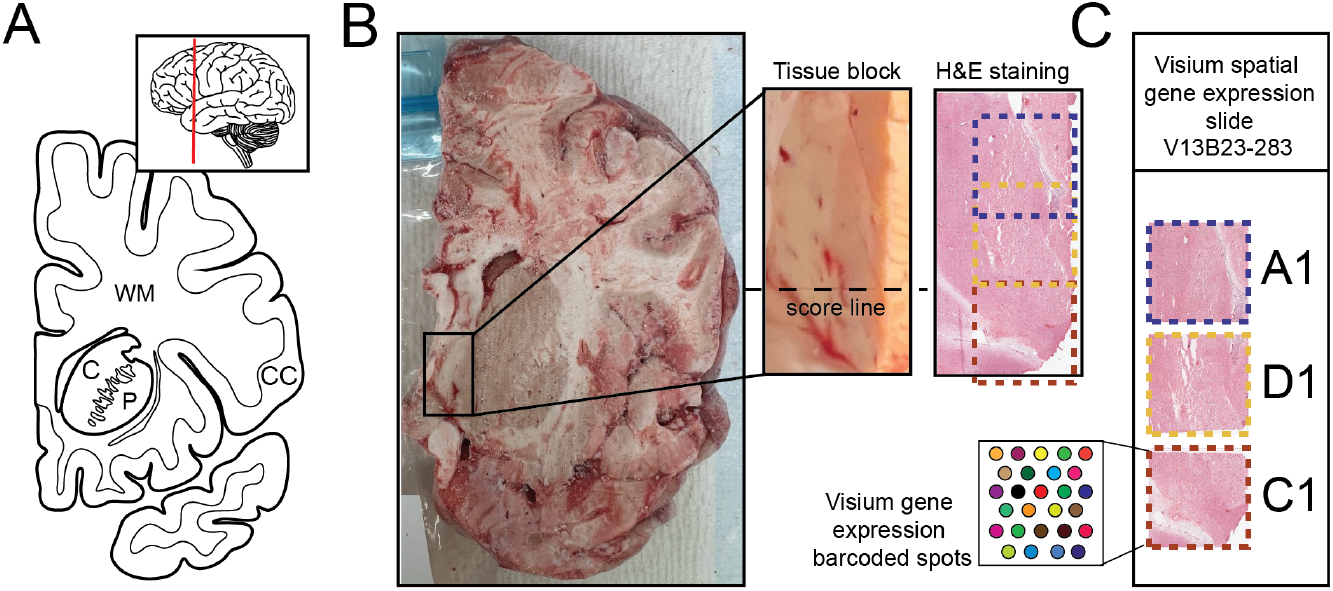
Experimental design to generate spatially-resolved transcriptomics (SRT) data from postmortem human brain across three Visium capture arrays. **(A)** Schematics illustrating a coronal human brain hemisphere at the level of the anterior striatum. Inset showing an illustration of the lateral side of the brain with a vertical red line depicting the location of the coronal slab. CC – cerebral cortex, C – caudate nucleus, P – putamen, WM – white matter. **(B)** Raw coronal brain slab, dissected tissue block with an indicated score line, and H&E staining of the 10 μm section taken from the same block. **(C)** A diagram depicting how tissue sections were arranged on the Visium slide with capture arrays A1 (blue) and D1 (yellow) arranged partially overlapping, while capture arrays D1 (yellow) and C1 (red) arranged adjacent to each other across a score line.

## Methods

### Post-mortem human tissue samples and tissue processing

Postmortem human brain tissue was obtained at the time of autopsy with informed consent from the legal next of kin, through the Department of Pathology, University of North Dakota School of Medicine and Health Sciences, under the WCG protocol #20111080. The brain donor was a 36-year-old female of European ancestry who died by natural causes (cardiac arrest), with no prior history of psychiatric, substance use, neurocognitive, or neurodegenerative disorders. The donor had a postmortem interval of 25 hours and a screening RNA integrity number (RIN) of 7.8 in prefrontal cortex. Details of tissue acquisition, handling, processing, dissection, clinical characterization, diagnoses, neuropathological examinations, and quality control measures have been described previously [12]. A tissue block of approximately 10 × 20 mm was dissected under visual guidance with a hand-held dental drill from a fresh frozen slab, encompassing the area medial to the anterior striatum. Following dissection, the tissue block was stored in a sealed cryogenic bag at -80°C. Prior to cryosectioning for the Visium assay, the tissue block was equilibrated to -20°C.

### Visium Data Generation

The block encompassing brain tissue medial to the anterior striatum was scored with a razor blade perpendicular to the boundary between the gray and white matter, and three 10μm sections were collected onto individual capture areas of the 10x Visium Gene Expression slide (part number 2000233, 10x Genomics). Tissue sections mounted onto two of the Visium capture areas were partially overlapping, while the third tissue sample was collected across the score mark, making it adjacent to one of the other sections, but not overlapping. Samples were processed according to manufacturer’s instructions (10x Visium Gene Expression protocol CG000239 rev D) as previously described [13]. In brief, the workflow includes tissue staining with hematoxylin and eosin, followed by high resolution image acquisition using an Aperio CS2 slide scanner (Leica) equipped with a 20x/0.75NA objective and 2x doubler. Following removal of the coverslip, tissue was permeabilized (18 minutes) to allow access to mRNA, followed by on-slide reverse transcription, collection of cDNA from the slide, and library construction. Libraries were quality controlled and sequenced on an Illumina NovaSeq X by Psomagen (Rockville, MD) according to manufacturer’s instructions at a minimum depth of 60,000 reads per Visium spot. Samples were sequenced to a median depth of 363,115,439 reads, corresponding to a median 82,209 of mean reads per spot, a median 1,158 of median unique molecular identifiers (UMIs) per spot, and median of 772 median genes per spot.

### Visium Data Processing

The high-resolution images obtained on the 10x Genomics Visium platform were processed using *VistoSeg* [14], a *MATLAB* -based pipeline that integrates gene expression data with histological data. First, the software splits the full-resolution image of the entire slide into individual capture areas (.tif file format), which are used as input to 1) *Loupe Browser* (10x Genomics) for fiducial frame alignment, and 2) *SpaceRanger* v3.0.0 to extract the spot metrics/coordinates [15]. FASTQ and image data were pre-processed with the 10x *SpaceRanger* pipeline version 3.0.0 [15]. The *Fiji* [8] instance of *ImageJ* v2.14.0 downloaded from https://imagej.net/software/fiji/downloads was used to interactively align images of each capture area. `build_spe()` from *visiumStitched* v0.99.0 was used to construct a *SpatialExperiment* 1.14.0 [9] object for downstream analysis and compatible with Bioconductor version 3.19. Data were filtered to remove genes that were not detected and spots that were out of tissue or had zero counts. Batch correction was not performed, as all data were from the same Visium slide. We performed log normalization by deconvolution using *scran* 1.32.0 [16] and *scuttle* 1.14.0 [17]. Spatially variable genes (SVGs) were found with *nnSVG* after filtering to genes with at least three counts in 0.5% of spots [18]. Clustering was performed using *PRECAST* v1.6.5 [11], using the top 500 SVGs by `rank` as input. Values of the additional gene-filtering parameters `premin.spots` and `postmin.spots` to `CreatePRECASTObject()` were set to 0 to retain all 500 input SVGs. To simulate analysis on the data without stitching, *nnSVG* was also run one capture area at a time, with the top 500 SVGs selected by averaging `rank` across capture areas before running *PRECAST*.

## Implementation

A *visiumStitched* workflow involves preparing the input Visium data to *Fiji*, before using R functions implemented in *visiumStitched* to build the stitched *SpatialExperiment* object (**Figure 2**).

**Figure 2.**
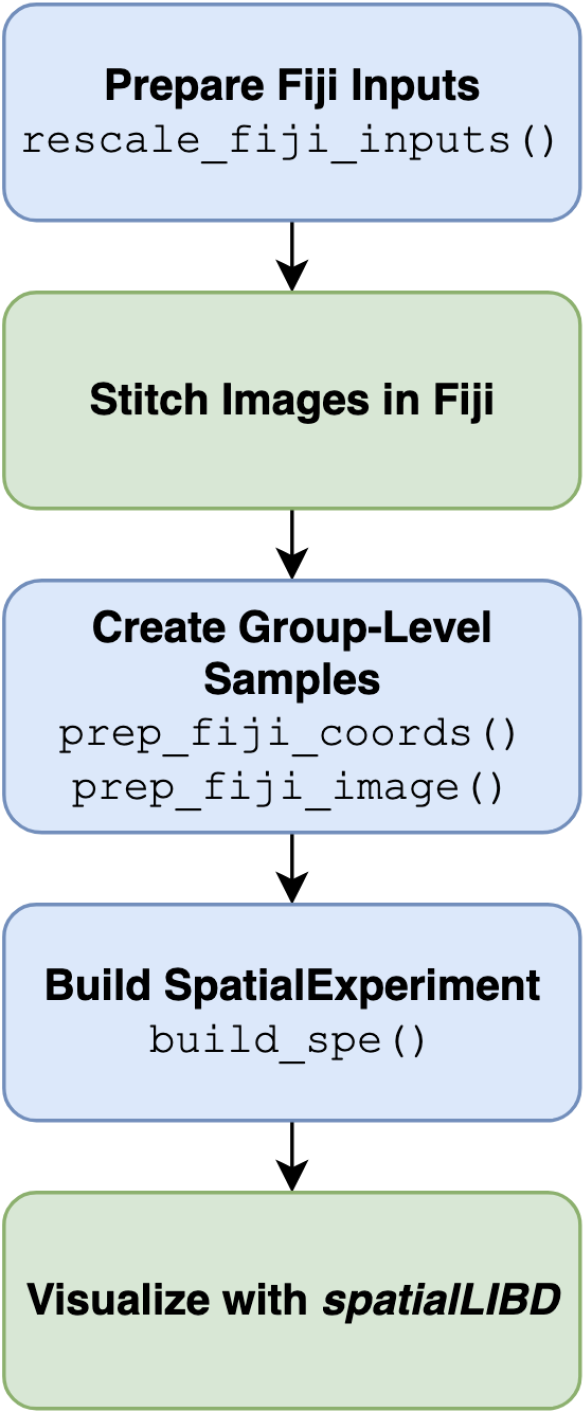
visiumStitched workflow. *visiumStitched* begins by scaling *SpaceRanger* [15] images to represent the same distance per pixel within each capture area group. Next, images for each group are stitched using *Fiji* [8]. The transformation specified by *Fiji* is used to create transformed spatial coordinates, and the stitched composite image is resized. Next, a *SpatialExperiment* [9] is constructed with one sample per group. The resulting stitched *SpatialExperiment* object can then be visualized with *spatialLIBD* [19]. Blue boxes represent functions implemented in *visiumStitched*, while green boxes are performed with external software.

### Preparing Inputs to *Fiji*

*SpaceRanger* [15] is typically used to process Visium data. As noted at https://www.10xgenomics.com/support/software/space-ranger/latest/analysis/outputs/spatial-outputs, `spaceranger --count` creates for each capture area a high resolution image (`tissue_hires_image.png`, up to 2,000 pixels in the longest dimension) and a file with rescaling factors linking the input user-supplied image and the images created by *SpaceRanger* (`scalefactors_json.json`). Note that we are using `tissue hires image.png` images given that *Fiji* cannot open full resolution images, which can vary in size depending on the acquisition parameters. For example, in a previous dataset the images had a median size of 2.3 GB each and 40x magnification [13]. The `tissue_hires_image.png` images are detailed enough for visualization purposes, and any image-based analysis pipeline that requires full resolution images would need to be applied before using *visiumStitched*, such as identifying number of cells per spot with *VistoSeg* [14].

In order to sensibly integrate several images with *Fiji*, they must be on the same scale; that is, a single pixel from each capture area to stitch must represent the same physical distance. To ensure this assumption is met given the nonuniformity of `tissue_hires_image.png` images, *visiumStitched*’s `rescale_fiji_inputs()` function rescales the `tissue_hires_image.png` image for each capture area *i* in a given group by the factor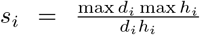. Here *d*_*i*_ is the number of pixels for one spot diameter and *h*_*i*_ is the high-resolution scale factor; these are given by the `spot_diameter_fullres` and `tissue_hires_scalef` values from *SpaceRanger*’s `scalefactors_json.json` file, respectively. The resulting image from `rescale_fiji_inputs(`*hires*_*i*_) will have dimensions equal to 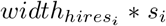 and 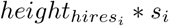.

### Stitching images with *Fiji*

Next, for each group of capture areas that should be stitched together, *Fiji* is run using the uniformly-scaled images produced by `rescale_fiji_inputs()`. With *Fiji*, the user manually stitches the images using manually defined tissue landmarks or prior knowledge (**Figure 3**). Manual image stitching is required for supporting stitching capture areas obtained across a score line which creates an actual physical gap, and thus cannot be done automatically. This feature expands the possible study design scenarios that *visiumStitched* supports.

**Figure 3.**
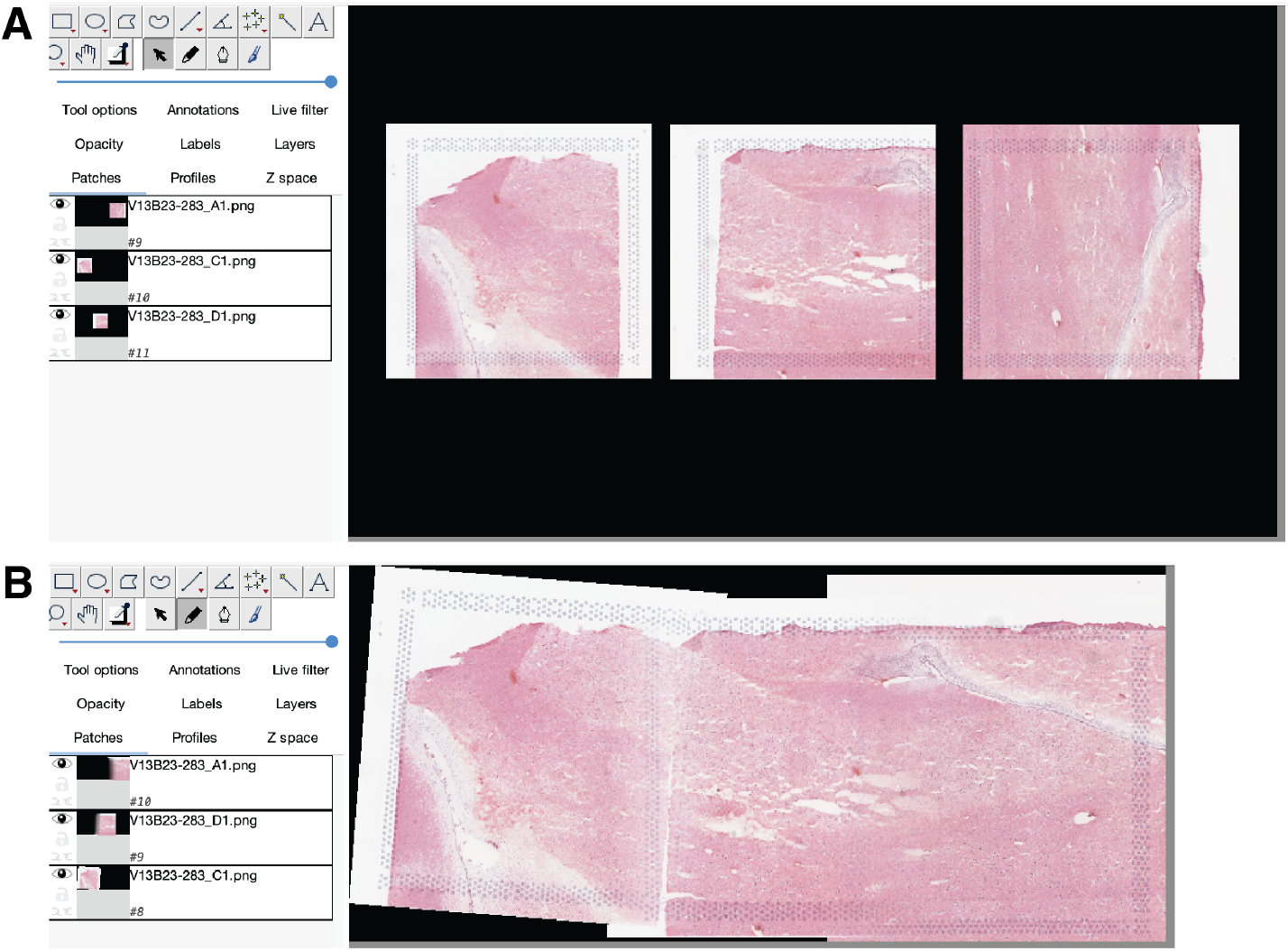
Stitching images in Fiji. **(A)** The three images for capture areas *V13B23-283_A1, V13B23-283_C1*, and *V13B23-283_D1* are imported in their original orientations from *SpaceRanger* onto a common canvas in *Fiji*. **(B)** Images are interactively rotated, translated, and blended to produce a stitched image for later use in the final stitched *SpatialExperiment* object.

For each group of capture areas that were stitched together, *Fiji* produces a high-resolution stitched image and XML file specifying a rigid affine transform per capture area. The user then constructs a table called `sample_info` specifying which capture areas were stitched together for each group.

### Creating Group-Level Samples

The `prep_fiji_coords()` function reads in the `tissue_positions.csv` *SpaceRanger* file for each capture area to obtain the original pixel coordinates, namely `pxl_row_in_fullres` and `pxl_col_in_fullres`. Next, the *Fiji* XML file is parsed for each group, and pixel coordinates for each capture area are multiplied by the specified matrix to apply the appropriate rigid affine transform. Finally, the coordinates are multiplied by a capture-area-specific scalar from `rescale_fiji_inputs()` called `intra_group_scalar`. Pixel coordinates are positions in terms of numbers of pixels, but the distance per pixel is not guaranteed to be precisely identical across capture areas in a group, particularly if different Visium slides are used. The scaling process ensures distances per pixel are consistent across all capture areas in a group. A new `tissue_positions.csv` is written for each group of stitched captured areas, containing the transformed pixel coordinates of all constituent capture areas.

The `prep_fiji_image()` function first resizes the stitched composite image from *Fiji* for each group to a default of 1,200 pixels in its longest dimension, assuming a square group composed of 2 by 2 stitched capture areas. The number of pixels in the longest dimension is configurable to handle different group compositions. This ensures each sample in the downstream *SpatialExperiment* is equal in resolution and memory usage to *SpaceRanger*’s `tissue_lowres_image.png` (up to 600 pixels in the longest dimension), as would typically be the case in a stitching-free experiment. The resized group image is written to a file called `tissue_lowres_image.png` with its compatible companion `scalefactors_json.json` file, which relates the pixels from the `lowres` stitched composite image to pixels in the `fullres` images just, like the files produced by *SpaceRanger*. Together the `prep_fiji_coords()` and `prep_fiji_image()` produce a directory imitating *SpaceRanger*’s spatial outputs, which can read into R using `build_spe()`.

### Building a *SpatialExperiment*

First, the `build_spe()` function from *visiumStitched* internally invokes `read10xVisiumWrapper()` from *spatialLIBD* [19] to build a *SpatialExperiment* [9] with one capture area per sample– just as may be done in an experiment without stitching. Next, the `imgData()` and `spatialCoords()` slots are overwritten with stitched images and pixel coordinates made with `prep_fiji_coords()` and `prep_fiji_image()`. The `sample_id` is overwritten to represent the group of stitched capture areas rather than each individual capture area. Note that all spots are preserved even in regions of overlap; an `exclude_overlapping_colData()` column is computed to indicate which spots to exclude from plots when using `spatialLIBD::vis_gene(is_stitched = TRUE)` or `spatialLIBD::vis_clus(is_stitched = TRUE)`. Finally, the array coordinates columns (`array_row` and `array_col` in `colData()`) are overwritten to be spatially contiguous within each group, as described next.

### Defining Array Coordinates for Stitched Samples

Following Visium’s hexagonal array design for a single capture area, `array_row` and `array_col` are columns in *SpaceRanger*’s `tissue_positions.csv` file representing each Visium spot’s row and column index as documented at https://www.10xgenomics.com/support/software/space-ranger/latest/analysis/outputs/spatial-outputs, that is, the array coordinates. By definition, `array_row` holds integer values from 0 to 77, while `array_col` holds integer values from 0 to 127. After performing image stitching, there is a need for new array coordinates relative to the new stitched image. The original array coordinates cannot reflect how capture areas may have been rotated and translated to reconstruct the true tissue arrangement across the capture areas (**Figure 4A**). To redefine these array coordinates, the function `add_array_coords()` assigns each group of capture areas a new Visium-like array, in place of the individual capture areas for that group. Like a true Visium array, artificial spots are arranged hexagonally with each a distance of 100 microns from its six neighbors; array rows and columns are defined such that the minimum and maximum values for `pxl_row_in_fullres` and `pxl_col_in_fullres` of any capture area for one donor correspond to values of 0 for `array_col` and `array_row`, respectively as in the original `tissue_positions.csv` file. Array rows and columns are added as necessary, generally well above the ordinary respective maximum values of 77 and 127, to cover the area spanned by all original spots for a whole stitched tissue section. Each original spot is then mapped to the nearest artificial spot by Euclidean distance to redefine the `array_row` and `array_col` values used downstream in spatial clustering via *BayesSpace* and *PRECAST* (**Figure 4B**) [10, 11]. Both these methods, in their default implementations for Visium data, use array coordinates for finding a spot’s neighbors, but are agnostic to the fact that more than one artificial spot may occupy the same array coordinates, as occurs in regions of overlap between capture areas (**Figure 4C**).

**Figure 4.**
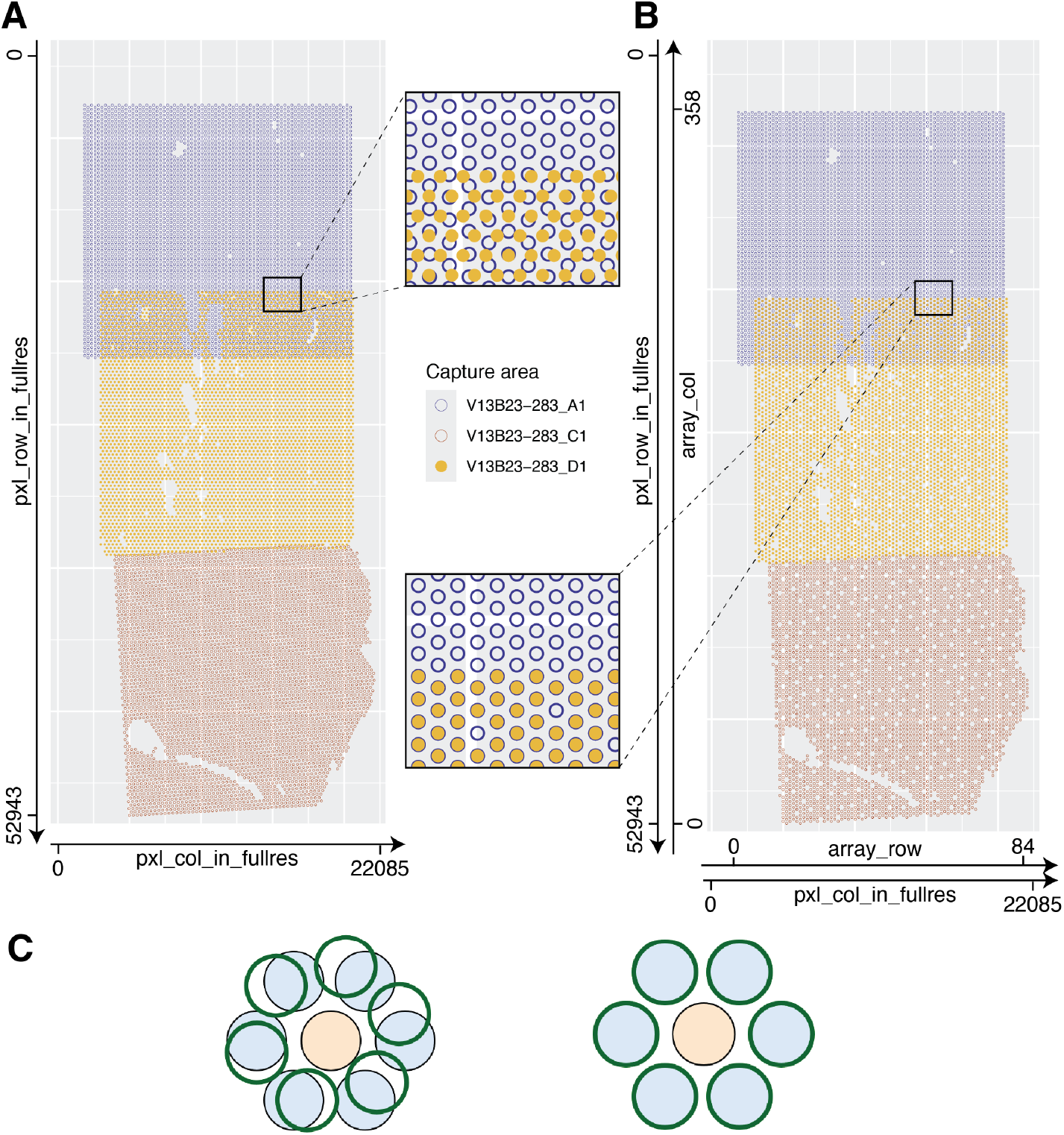
Defining array coordinates for stitched data. **(A)** The transformed pixel coordinates for donor Br2719 are plotted, colored by capture area. At the overlap of capture areas *V13B23-283_A1* and *V13B23-283_D1*, spot centroids do not cleanly overlap; the index of a given spot is poorly defined. **(B)** The transformed pixel coordinates for Br2719 are plotted, rounded to the nearest position on a hypothetical hexagonal, Visium-like array. At the overlap of capture areas *V13B23-283_A1* and *V13B23-283_D1*, spot centroids occupy the same exact positions. Array coordinates may be defined by indexing the row and column on the hypothetical Visium-like array encompassing the entire Br2719 sample. **(C)** Spots from two or more overlapping capture areas, filled in blue and outlined in green, respectively, can be assigned the same `array_col` and `array_row` values with *visiumStitched*, enabling using data from overlapping spots in downstream analyses that rely on the Visium hexagonal grid properties to find spot neighbors. On the left, exact spot positions are shown; on the right, positions rounded to the nearest array position are shown. The spot in orange has twelve unique neighbors at six unique array coordinates.

## Results

We have implemented the R/Bioconductor package *visiumStitched* that uniformly rescales images created by *SpaceRanger* that can then be manually stitched together with *Fiji* (**Figure 2**). *visiumStitched* then uses the *Fiji* and *SpaceRanger* outputs to build a *SpatialExperiment* R/Bioconductor object with precise pixel coordinates (**Figure 2**). We have also updated *spatialLIBD* to support visualization of *SpatialExperiment* objects created by *visiumStitched* (**Figure 2**). *visiumStitched* can also export *SpatialExperiment* objects for downstream analyses with *Seurat* [20].

To demonstrate the utility of *visiumStitched*, we generated example data from a postmortem human brain sample with three Visium capture areas (**Figure 1**) *spatialLIBD* (version 1.17.8 or newer) can be used to visualize stitched data, as showcased with excitatory neuron marker *SLC17A7* and the combined white matter markers *MBP*, GFAP, *PLP1*, and *AQP4* (**Figure 5A**). For overlapping spots *spatialLIBD* will only show spots from the capture area with higher mean UMIs, although data from all spots is contained in the stitched *SpatialExperiment* object produced by *visiumStitched*.

**Figure 5.**
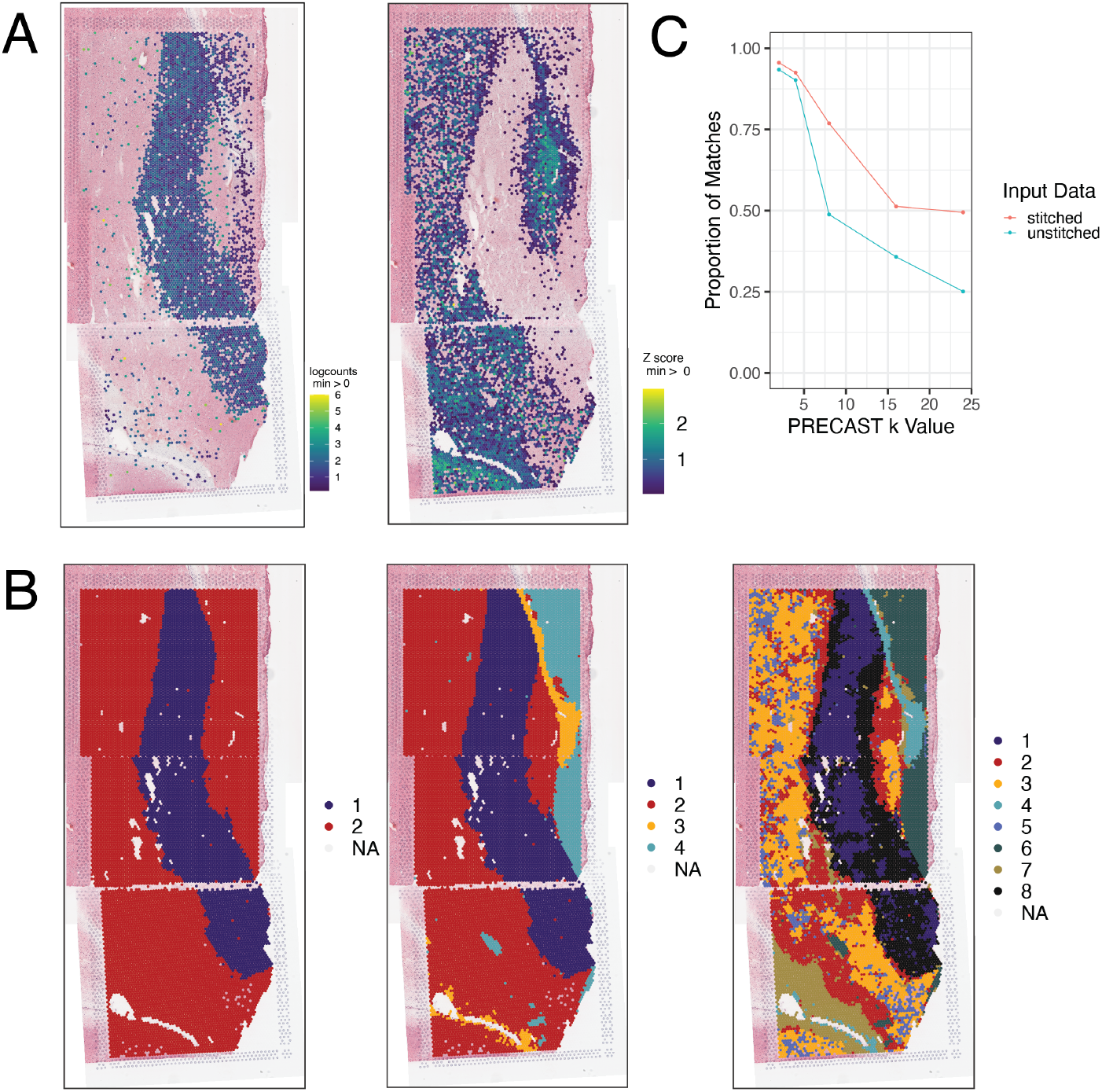
Gene expression and clustering of stitched data. **(A)** Expression of (left) *SLC17A7* and (right) the combination of white-matter marker genes *MBP, GFAP, PLP1, AQP4* is plotted for Br2719 using *spatialLIBD*. The white-matter marker genes were plotted using `multi_gene_method =’‘z_score’’` from `vis_gene()` implemented in *spatialLIBD*. **(B)** *PRECAST* cluster assignments are plotted for Br2719 at *k* = 2 and *k* = 4. NA spots were not assigned a cluster by *PRECAST*. **(C)** Proportion of agreement of cluster assignments at overlapping spots is plotted for *PRECAST* results at *k* = 2, *k* = 4, and *k* = 8. *PRECAST* tends to assign spots that overlap to the same clusters, but agreement declines as *k* increases. Stitching, rather than treating capture areas as separate samples, results in greater agreement.

*visiumStitched* constructs an artificial array hexagonal Visium grid to help downstream spatial clustering methods identify neighboring stitched spots based on a hexagonal grid pattern. Spatial clustering methods such as *BayesSpace* and *PRE-CAST* rely on the integer-based hexagonal grid to identify for each spot the six neighboring spot locations, yet they are compatible with having data from two or more spots on each of the six neighboring locations (**Figure 4C**). With the example human brain data, we identified 2, 4, and 8 spatially-resolved clusters using *PRECAST* (**Figure 5B**). Overlapping spots were frequently assigned to the same *PRECAST* clusters, although said frequency decreased as the number of clusters increased (**Figure 5C**). Stitching, compared with treating each capture area as a separate sample, results in greater agreement at overlaps; for `k = 8`, 49% of overlapping spots were assigned the same cluster without stitching, compared with 77% with stitching (a 57% relative increase).

An interactive website powered by *spatialLIBD* [19] at https://research.libd.org/visiumStitched_brain/ can be used to explore the stitched example human brain data.

## Discussion

*visiumStitched* provides a complete set of tools for stitching the gene-expression and imaging data from Visium experiments into analysis-ready R objects. We envision researchers primarily using *visiumStitched* for tissue regions of interest that are too large to fit onto a single Visium capture area.

The integration of separate images from spatial transcriptomics experiments is a well-established problem, and several solutions have been proposed [3, 4, 5, 6, 7]. However, to our knowledge, each solution proposed so far has one or more key limitations (**Table 1**). *STalign* [4] is a Python package that can align a pair of images, inferring an affine transform and optional diffeomorphic component that can also be applied to a target sample’s spatial coordinates. However, the source and target image must have considerable overlap, and therefore experiments with some adjacent or disjoint capture areas are not fully supported [4]. Moreover, for experiments that require stitching of three or more capture areas, users must write custom code to iteratively align pairs of capture areas, which becomes cumbersome for complex arrangements of many capture areas. In contrast, *visiumStitched* works with *Fiji*, which uses a graphical user interface to allow users to potentially align many capture areas which need not necessarily overlap. *Eggplant* and *GPSA* provide functionality to handle alignment of many capture areas, but are restricted to spatially adjacent replicates [5, 6] rather than partially overlapping capture areas. *PASTE2* may also handle many capture areas, but requires at least partial overlap among all capture areas [7], and thus cannot be used for adjacent tissue sections such as with D1 and C1 in the example data (**Figure 1B**). *VisiumStitcher* is closest to *visiumStitched* (**Table 1**), though *VisiumStitcher* does not create an artificial Visium array for downstream clustering analysis [3].

Given limitations of *Fiji, visiumStitched* works with the high resolution images (up to 2,000 pixels on the longest dimension) produced by *SpaceRanger*. This limitation does not affect visualization of the integrated image and gene expression data, but does limit the applicability of image-based analyses.

The core concepts of *visiumStitched* are extensible to handle newer versions of Visium by 10x Genomics. Visium v2 (Visium CytAssist) can support up to 11mm by 11mm capture areas and *visiumStitched* can process such data by changing the number of pixels on the longest dimension of the resulting stitched image. Visium HD, as well as the original Spatial Transcriptomics platform [21], use a square grid pattern of gene expression spots instead of a hexagonal grid. Methods such as *BayesSpace* [10] can perform spatially-aware clustering with square grid data by considering 8 neighboring spots. In the future, *visiumStitched*’s artificial hexagonal grid can be extended to construct artificial square grids, or other grid arrangements supported by *SpaceRanger* and downstream clustering methods.

## Conclusions

*visiumStitched* builds upon the existing functionality of *Fiji* to provide a complete workflow for researchers to stitch together gene-expression and imaging data for Visium-based spatial transcriptomics experiments. *visiumStitched* will facilitate the analysis of large tissue regions of interest that extend beyond a single Visium capture area by allowing researchers to treat the stitched region as a single continuous sample for downstream analysis. Since array coordinates are defined to be compatible with *SpaceRanger*’s array coordinates, the stitched *SpatialExperiment* [9] object is compatible with spatially aware clustering algorithms including *BayesSpace* [10] and *PRECAST* [11], enabling clustering to make use of the full spatial context of a large tissue region.

## Availability and Requirements

Project name: *visiumStitched*

Project home page: http://research.libd.org/visiumStitched/

Operating system(s): Windows, macOS, Linux

Programming language: R

Other requirements: *Fiji*

License: Artistic-2.0

Any restrictions to use by non-academics: N/A

## Declarations

Ethics approval and consent to participate Not applicable.

Consent for publication

Not applicable.

Availability of data and materials

The *visiumStitched* software is available from GitHub at https://github.com/LieberInstitute/visiumStitched [22], with documentation at http://research.libd.org/visiumStitched/. The analysis demonstrating the software’s usage on example brain data is available from GitHub at https://github.com/LieberInstitute/visiumStitched_brain [23]. The example human brain processed data is available through *spatialLIBD*’s `fetch_data()` function using version 1.17.8 or newer [19]. The raw data is available from the Globus endpoint ‘jhpce#visiumStitched brain’ that is also listed at http://research.libd.org/globus.

## Authors’ contributions

NJE, PR, and MT explored existing software, NJE and LCT designed *visiumStitched*, NJE implemented it, and MT beta-tested it. TMH dissected the human brain sample. SVB and YD generated the example Visium human brain data, and RAM processed the data. NJE, SVB, and LCT wrote the manuscript draft. SCP and KM supervised the data generation and project management. All authors contributed to revising the manuscript and approved the final manuscript.

## Acknowledgements

We thank the LIBD neuropathology team, particularly James Tooke and Amy Deep-Soboslay, for curation of the brain sample and assistance with tissue dissections. We thank Sarah Maguire for assistance with coordinating data sequencing requests with Psomagen. We thank the staff and physicians at the brain donation sites, and the generosity of the brain donors and their families, without whom this work would not be possible. We would like to thank the Joint High Performance Computing Exchange (JHPCE) for providing computing resources for these analyses. Finally, we thank the families of Connie and Steve Lieber and Milton and Tamar Maltz for their generous support.

## Abbreviations

Fiji: Fiji Is Just *ImageJ*
SPE: SpatialExperiment
UMI: Unique Molecular Identifier XML Extensible Markup Language SVG Spatially Variable Gene

